# Distinct tuning properties of human hippocampal neurons along the longitudinal axis during working memory

**DOI:** 10.64898/2026.09.03.749046

**Authors:** Morteza Mooziri, Ali Samii Moghaddam, Zahra Bahmani

**Affiliations:** Department of Brain and Cognitive Sciences, Cell Science Research Center, Royan Institute for Stem Cell Biology and Technology, ACECR, Tehran, Iran; School of Medicine, Zahedan University of Medical Sciences, Zahedan, Iran; Department of Electrical & Computer Engineering, Tarbiat Modares University, Tehran, Iran; Department of Cognitive Neuroscience, Faculty of Interdisciplinary Sciences and Technologies, Tarbiat Modares University, Tehran, Iran

## Abstract

Working memory (WM) is among the most sophisticated and fundamental capabilities of the mammalian brain. While the roles of prefrontal and sensory areas are heavily explored, there is little knowledge on how the hippocampus (HPC) contributes to this process. Here, we studied human HPC neuronal activities during a verbal WM task and reveal that neurons in the posterior HPC (PH) show more robust rate-modulations during WM. On the other hand, anterior HPC (AH) neurons are more prominently modulated by the phase of local and frontal cortex θ and αβ oscillations, a phenomenon that is accompanied by enhanced phase-synchronization between frontal cortex and HPC. Moreover, absence of correlational correspondence suggested that rate and phase are independent coding mechanisms. These results open a window to explore the functional dissociations along the primate HPC antero-posterior axis, a phenomenon long known to exist in the rodent brain. Furthermore, we suggest that combination of a variety of coding mechanisms in the human HPC neuronal population supports execution of WM.

**Graphical abstract:** 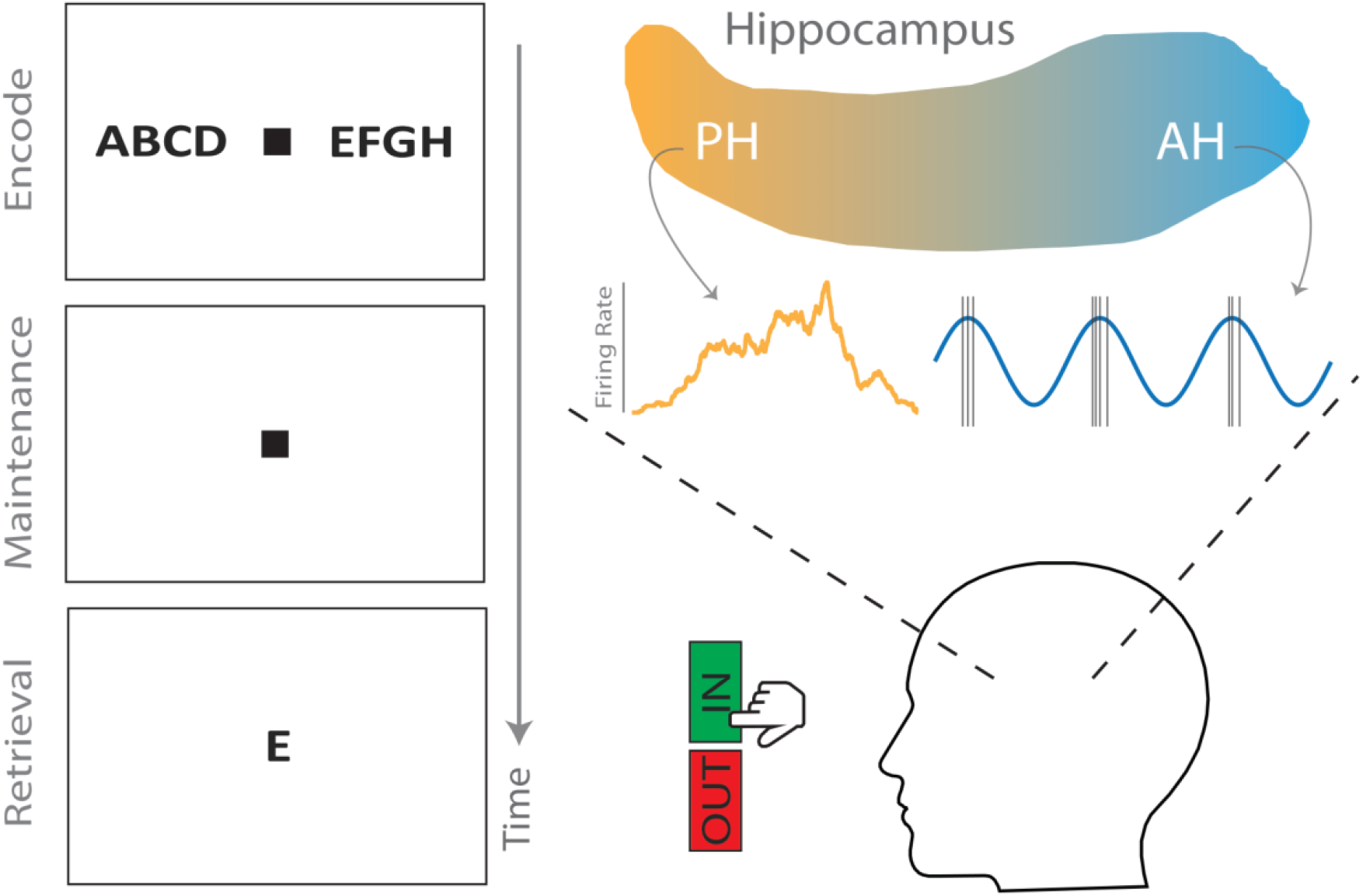

## Introduction

Working memory (WM) is a core cognitive process in the mammalian brain^1-6^. Through maintaining task-relevant information in the absence of sensory inputs, WM has fundamental roles in complex goal-directed behaviors, such as decision-making and learning^1-6^. Yet, despite significance, its neural underpinnings are largely unknown.

Execution of WM relies on coordinated activities between diverse brain regions, among which the roles of prefrontal (PFC) and sensory cortices are more prominent^1-8^. Traditionally, sustained neuronal firing in the PFC is believed to underlie WM, which is often termed as rate-coding regime^1-4,6-8^. On the other hand, tuning of PFC and/or sensory cortical neuronal spikes by local or distant slow oscillations, such as theta or alpha/beta bands, is reported to maintain WM content, known as phase-coding^3,9-13^. Especially in the latter case, PFC slow oscillations modulate neuronal activities in different brain regions during WM and attention^3,9-12^.

Recently, there has been a growing body of evidence introducing the hippocampus (HPC) as a neural substrate to support the maintenance of information^8,14-19^. In this line, both rate-^8,14-19^ and phase-^16,17^coding, acting as unrelated parallel processing mechanisms^17^, convey WM information in the HPC. On the other hand, rodent HPC is reported to be functionally distinct along the longitudinal axis, with the dorsal/ventral HPC playing a key role in cognitive/emotional processing^20,21^. However, despite sparse evidence^22-24^, there is lack of knowledge on how different poles of the human HPC contribute to different behaviors.

Here, we investigated human anterior (AH) and posterior (PH) HPC neuronal behaviors during verbal WM. We report that, during both encoding and maintenance, rate-modulation is more robust in the PH; on the other hand, tuning to local theta (θ; 1-10 Hz) and alpha/beta (αβ; 10-30 Hz) activities is stronger in AH during both task states. Furthermore, HPC neurons are phase-locked to frontal cortex θ and αβ oscillations, greater in AH, which we suggest it is due to the phase coupling between the oscillations in the two regions. Also, we show that, as suggested earlier^17^, the two coding regimes are separate mechanisms, and most probably, carry distinct WM-related information.

## Results

### Neurophysiology and behavior

We used a publicly available dataset of human neurophysiology during a verbal WM task^25^. The dataset comprised of Electroencephalography (EEG) as well as intracranial EEG (iEEG), which will be referred to as local filed potential (LFP) throughout the text, and neuronal spiking activities of human medial temporal lobe (MTL) sites during a modified Sternberg verbal WM task, in which series of English letters were used as visual stimuli^25^ (Fig. 1A). In each trial, after presentation of the stimuli (encoding section of the trial), the participants were supposed to hold the information in their memory (maintenance section of the trial) (Fig. 1A)^25^. During retrieval, subjects should have responded whether or not the presented item existed in the encoded items (Fig. 1A)^25^. The difficulty of different trials was manipulated by the number of encoded items; trials with four items (load 1) would be easier than six- and eight-item trials (loads 2 and 3, respectively). Details of the subjects’ information and demographics as well as their behavioral performance can be found in the main publications^14,25^.

**Figure 1.**
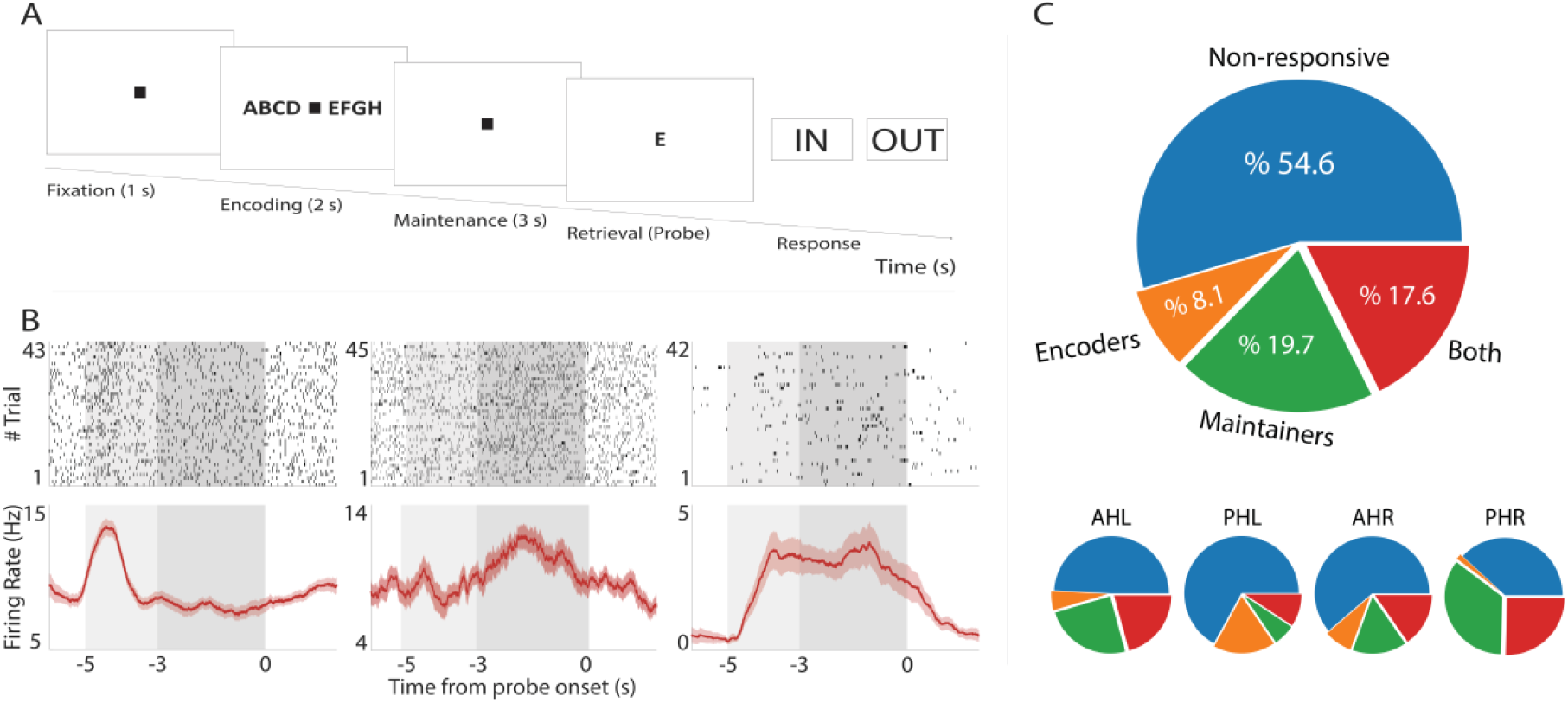
Task and general description of the HPC population. (A) Schematic illustration of the modified Sternberg verbal WM task. After fixation period (1 sec), a sample of 4,6 or 8 English letters appeared on the screen during the encoding (2 sec) section. Subsequently, the subject saw a blank screen during maintenance (3 sec) followed by presentation of a probe letter in the retrieval (2 sec). The subject should have responded whether or not, the probe letter existed in the encoded items.(B)Raster plots (upper panels) and PSTHs (lower panels) for a sample encoder (leftmost panels), maintainer (middle), and dual-functioning (rightmost panels) neurons. Light/dark rectangle denotes the encoding/maintenance section. In the PSTHs, solid line and shaded area indicate mean and SEM of firing rates, respectively. (C, upper) Pie chart describing the general functional structure of the HPC neuronal population. Assignment of each neuron to the corresponding category was decided based on the results of the two-sided Wilcoxon signed-rank test comparing encoding and maintenance with the fixation firing rates. (C, lower) Same as C, upper, but broken down for each sub-region and hemisphere. AHL, left anterior hippocampus; AHR, right anterior hippocampus; HPC, hippocampus; PHL, left posterior hippocampus; PHR, right posterior hippocampus; PSTH, peri-stimulus time histogram; WM, working memory.

HPC neurons maintain information in their firing rate for an immediate use^8,14-19,25^. Fig. 1B shows sample HPC neurons with selective activation during either the encoding (Fig. 1B, leftmost panels) or maintenance (Fig. 1B, middle panels) of information. Also, there are neurons that are responsive during both states; one sample of such neurons is illustrated in Fig. 1B, rightmost panels. Here, generally, we observed that 255/467 (%54.6) of the HPC neurons did not alter their firing in neither section of a trial, i.e., encoding or maintenance, compared to baseline, i.e., fixation period (non-responsive neurons; two-sided Wilcoxon signed-rank test, p > 0.05 for all cases; Fig. 1C, upper panel). 38/467 (%8.1) and 92/467 (%19.7) neurons showed a significant change in their firing rate only during encoding or maintenance (encoder or maintainer neurons, respectively; two-sided Wilcoxon signed-rank test with a p-value threshold of < 0.05 for all cases; Fig. 1C, upper panel); 82/467 (%17.6) neurons were categorized as both encoder and maintainer (two-sided Wilcoxon signed-rank test, p < 0.05 for each condition; Fig. 1C, upper panel). Also, we observed some degree of heterogeneity between different HPC sub-regions and hemispheres. Specifically, in the left anterior HPC (AHL), %49.2 of the population were non-responsive, %5.5 were encoders, %24.6 were maintainers, and %20.8 were both encoder and maintainer; these portions were %67.0,%17.5, %6.2, and %9.3, respectively, in left posterior HPC (PHL), %61.3, %8.1, %15.3, and %15.3, respectively, in right anterior HPC (AHR), and %38.1, %1.6, %34.9, and %25.4, respectively, in right posterior HPC (PHR). Overall, roughly half the neurons of human HPC are rate-modulated in at least one stage of a WM process.

Subjects of this dataset have attended more than one recording session^25^. Thus, hypothetically, some neurons might have been captured more than once. To exclude this effect on neuronal results, we repeated these analyses on a dataset of only each subject’s first session (9 sessions), the results of which will be described in the appropriate context. Here, the general functional structure of the HPC neuronal population is similar between the two datasets (Fig. 1A and Supplementary Fig. 1A, upper panels). However, we observed some degree of heterogeneity between the two datasets; for example, surprisingly, we did not observe any only-encoder neurons in right HPC in the smaller dataset (Supplementary Fig. 1A, lower panels).

### WM rate modulation is stronger in PH

A substantial body of literature shows longitudinal differentiation along the HPC in different species^20-24,26,27^. This issue is more heavily studied in the rodent brain, where the ventral (vHPC) and dorsal (dHPC) sub-regions are linked to distinct behaviors^20,21^. While the rodent dHPC, which corresponds to the primate posterior HPC (PH), plays a key role in cognitive functions like memory and spatial navigation, the rodent vHPC, which corresponds to the primate anterior HPC (AH), predominantly engages in processing of emotions like fear and anxiety^20,21,26,28^. Also, there is evidence, yet sparse and inconclusive, on such functional specialization along the human HPC longitudinal axis^22-24^. For instance, human PH dynamically modulates the AH activity during WM^24^. With this perspective, we tried to see whether or not the human AH and PH contribute differently to WM processing. Therefore, to make more concrete inferences on rate modulation, we computed the encode and maintenance d-prime (d’; see Methods, Statistical analyses) of every unit. For that, each neuron’s encode (encode d’) and maintenance (maintenance d’) firing rates were separately compared to the same neuron’s baseline firing rate; this metric reflects the strength of activation for each neuron in each stage. Fig. 2A shows the encode and maintenance d’ for all neurons in the population, where the encode d’ ranged from -2.94 to 1.90 and the maintenance d’ from -1.02 to 1.45. We observed that %25.08 of AH neurons and %26.88 of PH neurons showed rate modulation during encoding; these portions were %39.41 and %33.12, respectively, for maintenance. Next, these populations of modulated neurons were used to compare the strength of activation between the two HPC poles. We found that both encode (mean ± SEM; d’_AH_: 0.50 ± 0.003, d’_PH_: 0.67 ± 0.01; n = 77 and 43 neurons, respectively; p = 0.04, Mann-Whitney test; Fig. 2B, left panel) and maintenance (mean ± SEM; d’_AH_: 0.49 ± 0.002, d’_PH_: 0.67 ± 0.006; n = 121 and 53 neurons, respectively; p = 0.0005, Mann-Whitney test; Fig. 2B, right panel) rate modulations are stronger in PH, compared to AH. Also, generally, similar results were observed in the sub-dataset; the encode d’ ranged from -2.41 to 1.09 and the maintenance d’ from -1.02 to 1.45. The proportion of activated neurons during encoding and maintenance were %30.68 and %40.91 in AH as well as %29.09 and %38.18 in PH, respectively. Also, while maintenance rate modulations were stronger in PH, compared to AH (mean ± SEM; d’_AH_: 0.50 ± 0.006, d’_PH_: 0.73 ± 0.01; n = 36 and 21 neurons, respectively; p = 0.003, Mann-Whitney test; Supplementary Fig. 1B, right panel), this effect did not reach significance during encoding (mean ± SEM; d’_AH_: 0.46 ± 0.007, d’_PH_: 0.69 ± 0.03; n = 27 and 16 neurons, respectively; p = 0.16, Mann-Whitney test; Supplementary Fig. 1B, left panel).

**Figure 2.**
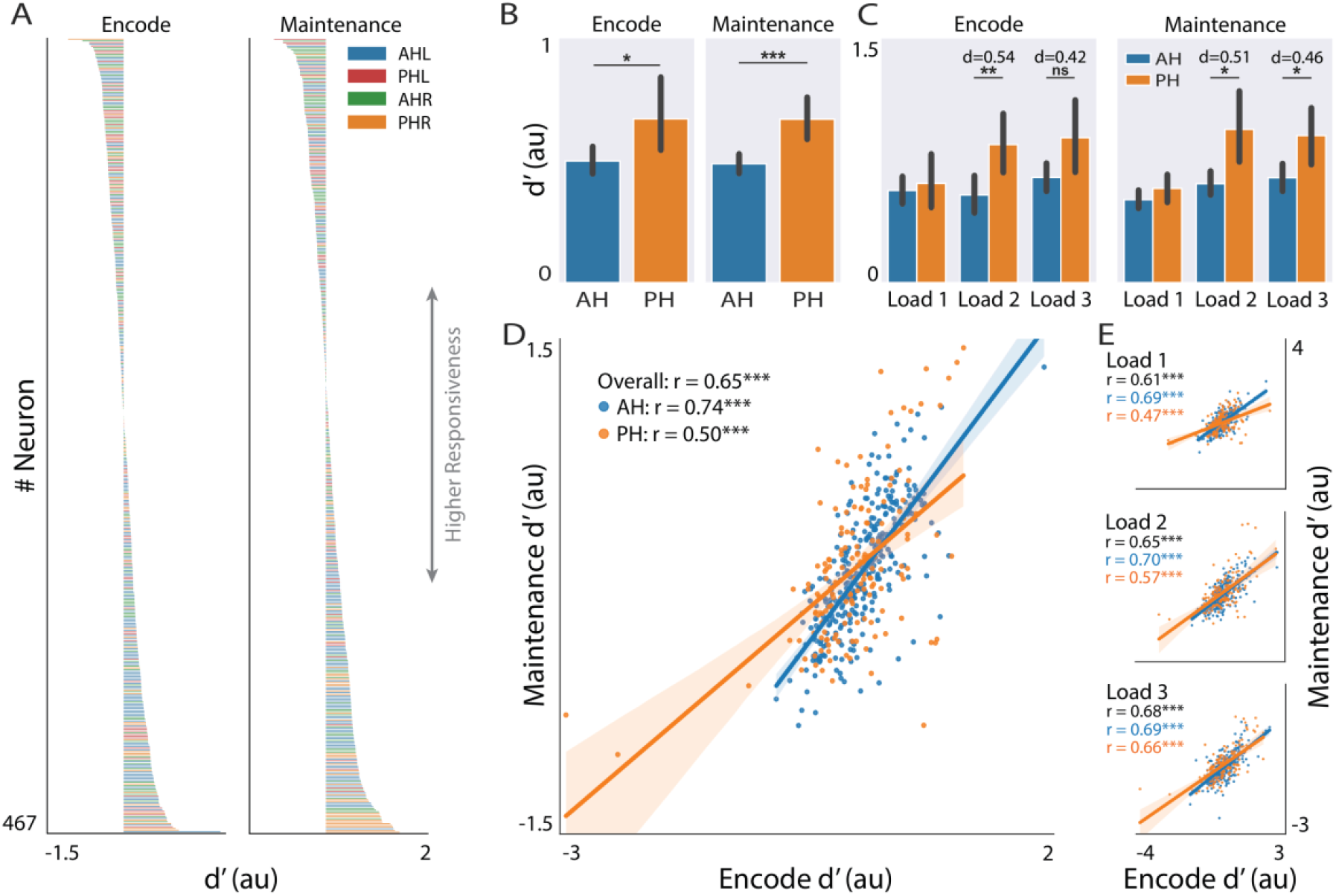
Rate modulation of the HPC neurons during WM. (A) Encode (left panel) and maintenance (right panel) d’ values for all HPC neurons (n=467). Colors represent region of neuron’s placement; two neurons with the most negative encode d’ values (d’_encode_ = -2.94 and d’_encode_ = -2.41) are excluded from visualization (available in analyses). (B) Statistical comparison of encode (left panel) and maintenance (right panel) d’ between rate modulated neurons in the AH (n_encode_ = 77, n_maintenance_ = 121; blue columns) and PH (n_encode_ = 43, n_maintenance_ = 53; orange columns). Bars and error bars represent mean and %95 CI, respectively. Mann-Whitney test. (C) Same as B, but for each load. d values indicate effect sizes based on Cohen’s d. (D) Correlation between encode d’ and maintenance d’. Each point is a neuron. Solid lines and shaded areas denote the fitted linear regression models and the %95 CI of the regression. Spearman correlation. (E) Same as D, but for each load. * p < 0.05, ** p < 0.01, *** p < 0.001. AH, anterior hippocampus; CI, confidence interval; HPC, hippocampus; PH, posterior hippocampus; WM, working memory.

Previous works suggest that human HPC neurons’ rate modulations might depend on the difficulty of the task^14,15^; generally, some HPC neurons are activated more in higher loads^14,15^. Thus, we sought to find out if the observed difference between AH and PH activation is stronger in higher loads. We observed that there is no difference between the two regions in load 1, which is the simplest type of trials, neither during encoding (mean ± SEM; d’_AH_: 0.57 ± 0.005, d’_PH_: 0.61 ± 0.01; n = 77 and 43 neurons, respectively; p = 0.76, Cohen’s d = 0.09, Mann-Whitney test; Fig. 2C, left panel) nor maintenance (mean ± SEM; d’_AH_: 0.51 ± 0.003, d’_PH_: 0.58 ± 0.006; n = 121 and 53 neurons, respectively; p = 0.16, Cohen’s d = 0.21, Mann-Whitney test; Fig. 2C, right panel). Interestingly, we observed that when the task got harder, the PH-AH rate modulation difference became more prominent, i.e., larger effect sizes. Specifically, this difference was evident in load 2 for both encoding (mean ± SEM; d’_AH_: 0.54 ± 0.007, d’_PH_: 0.85 ± 0.02; n = 77 and 43 neurons, respectively; p = 0.003, Cohen’s d = 0.54, Mann-Whitney test; Fig. 2C, left panel) and maintenance (mean ± SEM; d’_AH_: 0.61 ± 0.004, d’_PH_: 0.94 ± 0.02; n = 121 and 53 neurons, respectively; p = 0.02, Cohen’s d = 0.51, Mann-Whitney test; Fig. 2C, right panel), and in load 3 during maintenance (mean ± SEM; d’_AH_: 0.64 ± 0.004, d’_PH_: 0.90 ± 0.01; n = 121 and 53 neurons, respectively; p = 0.01, Cohen’s d = 0.46, Mann-Whitney test; Fig. 2C, right panel). For encoding of load 3 trials, we observed a moderate, yet insignificant, effect (mean ± SEM; d’_AH_: 0.65 ± 0.005, d’_PH_: 0.89 ± 0.02; n = 77 and 43 neurons, respectively; p = 0.11, Cohen’s d = 0.42, Mann-Whitney test; Fig. 2C, left panel). Together, these results show that the human PH neurons are more robustly rate-modulated to hold the content of WM.

Furthermore, we asked whether or not the level of activation of HPC neurons in the encoding section was predictive of their activation in the maintenance section, and vice versa. We observed a positive correlation between encode d’ and maintenance d’ (total: r = 0.65, p < 1e-56; spearman correlation; Fig. 2D). Moreover, this effect holds true in each region (AH: r = 0.74, p < 1e-53; PH: r = 0.50, p < 1e-10; spearman correlation; Fig. 2D) and each load (load 1; total: r = 0.61, p < 1e-47; AH: r = 0.69, p < 1e-42; PH: r = 0.47, p < 1e-9; load 2; total: r = 0.65, p < 1e-56; AH: r = 0.70, p < 1e-44; PH: r = 0.57, p < 1e-14; load 3; total: r = 0.68, p < 1e-63; AH: r = 0.69, p < 1e-44; PH: r = 0.66, p < 1e-20; spearman correlation; Fig. 2E). We also observed the same results in the sub-dataset (statistics are provided within the figure, Spearman correlation; Supplementary Fig. 1C,D). To rule-out the possibility that this correlation is driven by non-rate-modulated neurons, we repeated the same procedure with either encoder (total: r = 0.67, p < 1e-15; AH: r = 0.69, p < 1e-11; PH: r = 0.60, p < 1e-4; spearman correlation; Supplementary Fig. 1E, upper panel) or maintainer (total: r = 0.62, p < 1e-18; AH: r = 0.70, p < 1e-18; PH: r = 0.59, p < 1e-5; spearman correlation; Supplementary Fig. 1E, lower panel) neurons, and found significant correlations in both cases. These results suggest that the degree of each neuron’s contribution to the maintenance of WM content is related to its activation while the HPC is receiving the information.

### WM phase modulation is stronger in AH

Besides rate modulation, many human HPC neurons are tuned to fire at a specific phase of the local low frequency oscillations^16,17^. One sample of such neurons is depicted in Fig. 3A, which shows a clear tendency to fire at a particular phase of the 7 Hz oscillations during both encoding and maintenance. Therefore, next, we sought to compare the locking of neurons to regional oscillations between the two poles of HPC, for which we quantified each neuron’s spike-phase locking (SPL). Since there are several LFP recording sites (channels) in each patient’s HPC, we computed the locking of each neuron to all possible frequencies, from low θ to high β, of the oscillations recorded from all channels, as previously described^17^ (see Methods, Neural data analysis). Next, at every frequency, each neuron’s maximal shuffle-corrected SPL value (200 times shuffling across timepoints and trials) among all the channels was extracted (see Methods, Neural data analysis). Also, if a distribution of spike phases was uniform (insignificant Rayleigh test at a p-value level of 0.05), the corresponding SPL value was discarded (see Methods, Neural data analysis). The sample neuron illustrated in Fig. 3B has a median firing phase of ϕ_encode_ = 0.05 rad and phase locking strength of SPL_encode_ = 26.83 (raw SPL = 0.22) during encoding and median firing phase of ϕ_maintenance_ = 0.22 rad and phase locking strength of SPL_maintenance_ = 48.89 (raw SPL = 0.24) during maintenance, with respect to the local 7-Hz oscillation. Overall, we observed that 446/454 (%98.24) and 402/454 (%88.55) HPC neurons had significant locking to at least one channel-frequency pair during encoding in the θ and αβ bands, respectively. These numbers were 447/454 (%98.46) and 407/454 (%89.65) for maintenance, respectively. The median θ and αβ phase locking across this population of tuned neurons was SPL_θ_ = 5.03 and SPL_αβ_ = 4.15 during encoding and SPL_θ_ = 5.53 and SPL_αβ_ = 4.31 during maintenance, respectively. The preferred frequency (defined as the frequency with maximal SPL value; see Methods, Neural data analysis) differed among neurons, with an average preferred frequency of F_overall-encode_ = 8.41 Hz and F_overall-maintenance_ = 7.78 Hz in general (the entire range of studied frequencies, i.e., 1-30 Hz), F_θ-encode_ = 3.89 Hz and F_θ-maintenance_ = 3.89 Hz in the θ band, and F_αβ-encode_ = 17.18 Hz and F_αβ-maintenance_ = 17.32 Hz in the αβ band during encoding and maintenance, respectively.

**Figure 3.**
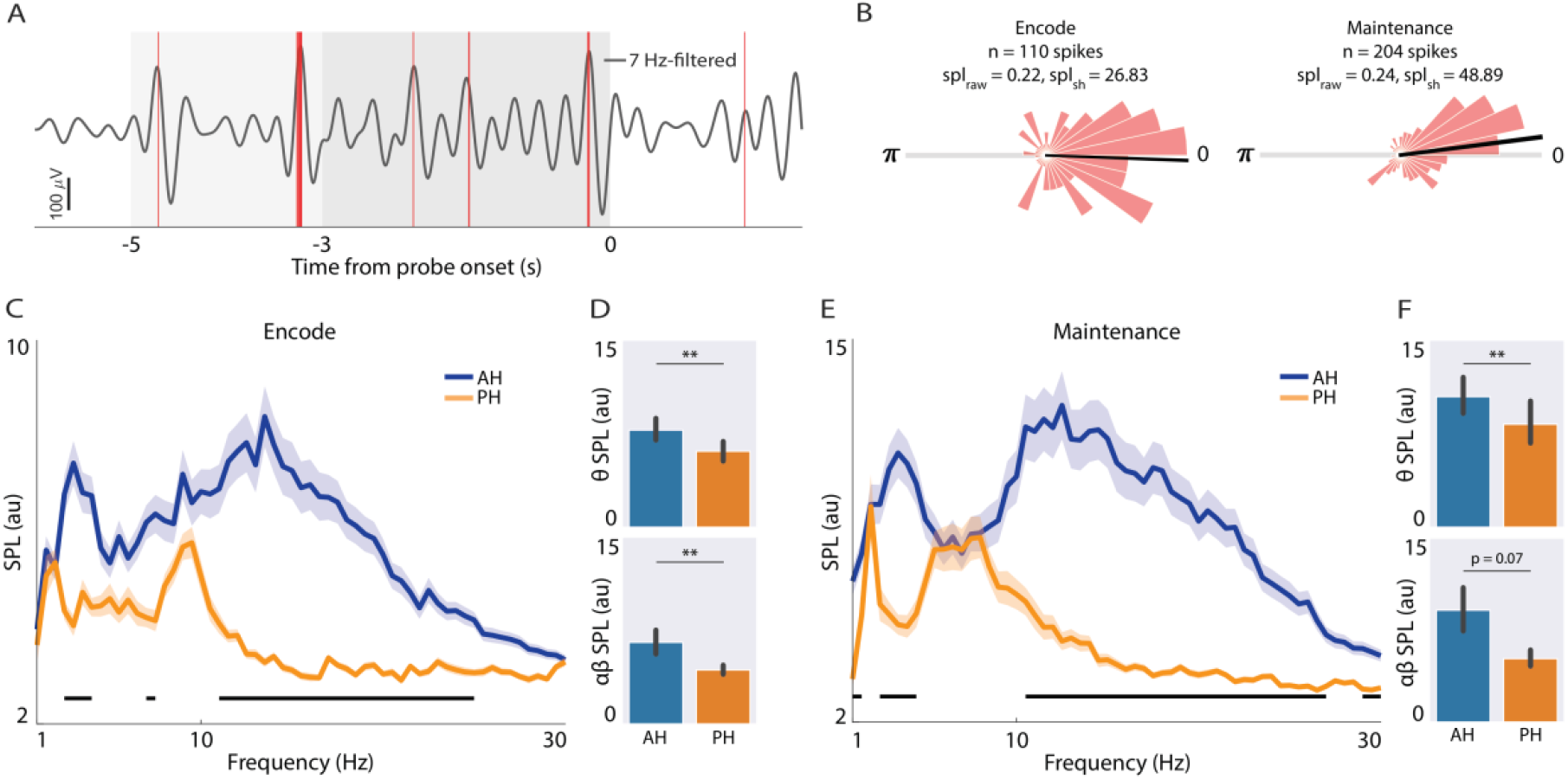
Phase modulation of the HPC neurons during WM. (A) A schematic of the temporal relationship between neuronal spikes and an oscillation. Sample for a 7-Hz filtered signal from a HPC site (black solid line) in a trial and the simultaneous spikes recorded from a neuron (red vertical lines) in the same trial. Light/dark rectangle denotes the encoding/maintenance section. (B) Polar histograms for all the spike-phases of a sample neuron (across all trials), with respect to the local 7-Hz oscillation, during encoding (left panel) and maintenance (right panel). Black lines denote the median phase of spike times, with respect the oscillation. (C,E) Shuffle-corrected local SPL values of AH and PH neurons. Solid lines and shaded areas indicate mean and SEM of SPL_sh_ values, respectively. Horizontal black lines in the bottom indicate frequencies with statistically different SPL_sh_ values (at p = 0.05) between AH and PH. Permutation test. (D,F) Statistical comparison of encode (D) and maintenance (F) SPL_sh_ between AH (n_θ-encode_ = 290, n_θ-maintenance_ = 291, n_αβ-encode_ = 266, n_αβ-maintenance_ = 270; blue columns) and PH (n_θ-encode_ = 156, n_θ-maintenance_ = 156, n_αβ-encode_ = 136, n_αβ-maintenance_ = 137; orange columns) neurons in the θ (upper panels) and αβ (lower panels) frequency bands. Bars and error bars represent mean and %95 CI, respectively. Mann-Whitney test. ** p < 0.01. AH, anterior hippocampus; CI, confidence interval; HPC, hippocampus; PH, posterior hippocampus; SPL_raw_/SPL_sh_, raw/shuffle-corrected spike-phase locking value; WM, working memory.

Next, we tested whether the phase-modulation of neurons was different between the two poles of the HPC. We found that AH neurons show stronger phase-tuning to both θ (mean ± SEM; SPL_AH_: 7.80 ± 0.03, SPL_PH_: 6.10 ± 0.03; n = 290 and 156 neurons, respectively; p = 0.004, Mann-Whitney test; Fig. 3C and Fig. 3D, upper panel) and αβ (mean ± SEM; SPL_AH_: 6.55 ± 0.03, SPL_PH_: 4.33 ± 0.02; n = 266 and 136 neurons, respectively; p = 0.005, Mann-Whitney test; Fig. 3C and Fig. 3D, lower panel) oscillations during encoding, compared to PH neurons. Also, AH neurons were more prominently coupled to θ frequency band during maintenance (mean ± SEM; SPL_AH_: 10.48 ± 0.05, SPL_PH_: 8.27 ± 0.07; n = 291 and 156 neurons, respectively; p = 0.001, Mann-Whitney test; Fig. 3E and Fig. 3F, upper panel), while this effect did not reach significance for αβ oscillations (mean ± SEM; SPL_AH_: 9.12 ± 0.06, SPL_PH_: 5.17 ± 0.03; n = 270 and 137 neurons, respectively; p = 0.07, Mann-Whitney test; Fig. 3E and Fig. 3F, lower panel). Similar results were observed in the sub-dataset (data not shown). Overall, these results suggest that while PH shows more prominent rate-modulation, AH neurons are primarily phase-modulated to maintain information during WM.

### Frontal cortex oscillations modulate the activity of HPC neurons during WM

Beyond the local oscillations, some HPC neurons are tuned to distal oscillations^16^. In this case, coupling of HPC neurons to frontal low frequency rhythms is an important coding mechanism of WM^16^. To further study this effect, we computed the phase locking of HPC neurons to frontal θ and αβ oscillations, for which we used the F3 and F4 electrodes of EEG (hemispheric relation was respected). We observed that 367/454 (%80.84) and 182/454 (%40.09) HPC neurons had significant locking to at least one frequency of the frontal EEG during encoding in the θ and αβ bands, respectively. These numbers were 368/454 (%81.06) and 178/454 (%39.21) for maintenance, respectively. The median θ and αβ phase locking across this population of tuned neurons was SPL_θ_ = 3.00 and SPL_αβ_ = 2.96 during encoding and SPL_θ_ = 3.15 and SPL_αβ_ = 3.15 during maintenance, respectively. The preferred frequency differed among neurons, with an average preferred frequency of F_overall-encode_ = 6.39 Hz and F_overall-maintenance_ = 6.46 Hz in general, F_θ-encode_ = 3.10 Hz and F_θ-maintenance_ = 2.88 Hz in the θ band, and F_αβ-encode_ = 13.31 Hz and F_αβ-maintenance_ = 13.52 Hz in the αβ band during encoding and maintenance, respectively.

Next, we tried to see whether or not this tuning differs between AH and PH. We found that during encoding, AH neurons were more strongly phase-tuned to both θ (mean ± SEM; SPL_frontal-AH_: 3.70 ± 0.01, SPL_frontal-PH_: 3.03 ± 0.02; n = 237 and 130 neurons, respectively; p = 0.02, Mann-Whitney test; Fig. 4A and Fig. 4B, upper panel) and αβ (mean ± SEM; SPL_frontal-AH_: 3.79 ± 0.01, SPL_frontal-PH_: 3.16 ± 0.03; n = 126 and 56 neurons, respectively; p = 0.0007, Mann-Whitney test; Fig. 4A and Fig. 4B, lower panel) rhythms of the frontal cortex, compared to PH. Also, AH neurons were more prominently locked to the frontal θ oscillations during maintenance, compared to the PH neurons (mean ± SEM; SPL_frontal-AH_: 3.97 ± 0.01, SPL_frontal-PH_: 3.34 ± 0.02; n = 242 and 126 neurons, respectively; p = 0.04, Mann-Whitney test; Fig. 4C and Fig. 4D, upper panel). However, we did not observe a significant difference between tuning of AH and PH neurons to frontal αβ (mean ± SEM; SPL_frontal-AH_: 3.55 ± 0.01, SPL_frontal-PH_: 3.40 ± 0.03; n = 126 and 52 neurons, respectively; p = 0.04, Mann-Whitney test; Fig. 4C and Fig. 4D, upper panel). Also, the sub-dataset showed similar results (data not shown). As previously shown, frontal θ rhythms play a key role in tuning the firing of HPC neurons while information is being held in WM^16^. Additionally, here we show that this tuning differs between the two poles of the HPC. These results, along with the above (Fig. 3), further suggest that phase-modulation of neuronal activities is more important in AH, compared to PH.

**Figure 4.**
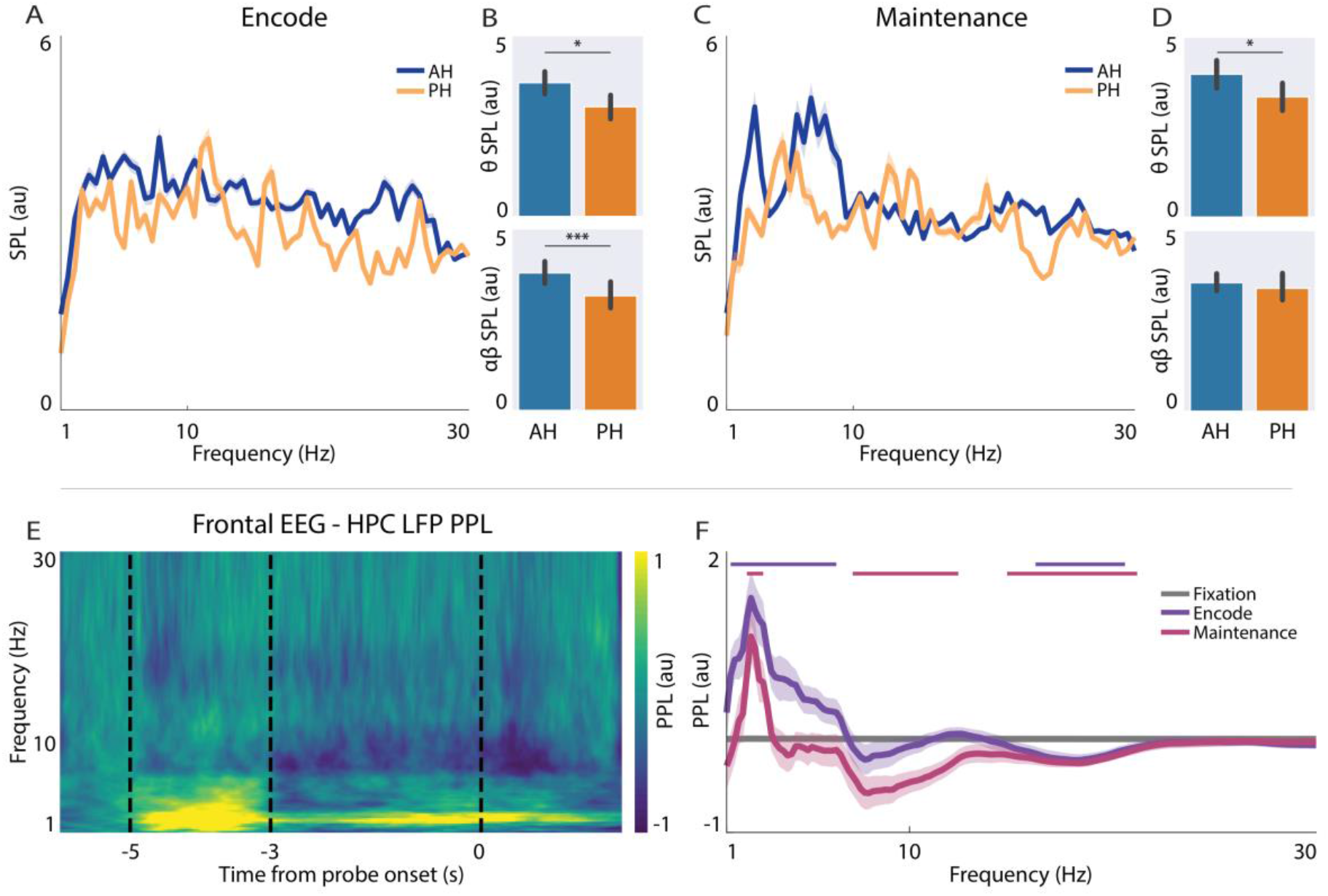
Frontal cortex tuning of HPC neuronal activity during WM. (A,C) Shuffle-corrected SPL values of AH and PH neurons with respect to frontal cortex EEG. Solid lines and shaded areas indicate mean and SEM of SPL values, respectively. (B,D) Statistical comparison of encode (D) and maintenance (F) SPL between AH (n_θ-encode_ = 237, n_θ-maintenance_ = 242, n_αβ-encode_ = 126, n_αβ-maintenance_ = 126; blue columns) and PH (n_θ-encode_ = 130, n_θ-maintenance_ = 126, n_αβ-encode_ = 56, n_αβ-maintenance_ = 52; orange columns) neurons in the θ (upper panels) and αβ (lower panels) frequency bands. Bars and error bars represent mean and %95 CI, respectively. Mann-Whitney test. (E) Heatmap indicating average baseline-corrected PPL between frontal EEG and simultaneous HPC LFP. (F) Time-averaged baseline corrected PPL between frontal EEG and simultaneous HPC LFP in encode, maintenance, and fixation periods. Solid lines and shaded areas indicate mean and SEM of PPL, respectively. Horizontal pink/purple line in the top indicates frequencies with statistically different PPL value (at p = 0.05) compared to baseline during encoding/maintenance. Permutation test. * p < 0.05, *** p < 0.001. AH, anterior hippocampus; CI, confidence interval; EEG, electroencephalography; HPC, hippocampus; PH, posterior hippocampus; SPL, spike-phase locking; WM, working memory.

Since the HPC subregion with stronger tuning to local oscillations, i.e., AH, was also the one with greater locking to frontal cortex rhythms, we were curious to see if there is a functional connectivity between the two regions’ (i.e., frontal cortex and HPC) oscillations. To address that, we computed the phase-phase locking (PPL; see Methods) between frontal EEG and HPC LFP. Interestingly, we found a prominent functional connectivity between frontal cortex and HPC θ oscillations during both encoding and maintenance (Fig. 4E,F; p-values computed through permutation; frequencies that showed a significant difference compared to baseline at the level of p = 0.05 are indicated with horizontal lines in Fig. 4F). The peak θ coupling occurred at F_encode_ = 2.2 Hz and F_maintenance_ = 2.2 Hz during encoding and maintenance,respectively. Together, these observations suggest that oscillatory mechanisms might convey the signal from frontal cortex to HPC, tuning the activity of HPC neurons, and keep the information in the HPC memory reservoir.

### Phase-rate interactions during WM in HPC

Phase- and rate-coding are distinct WM mechanisms in the human HPC^17^. Thus, next, we tried to check the relationship between phase and rate modulations. For that, we checked their correlational correspondence in our population and found that they were not correlated in most conditions (separated for modulating frequency band and neuron type), except for the following weak values: HPC θ (r = 0.25, p < 1e-4) and αβ (r = 0.20, p = 0.002) bands during maintenance in enhancive neurons. Next, we computed the pairwise correlation between all metrics (Supplementary Fig. 2A, left panel), and hypothesized that if the two coding mechanisms are unrelated, the data-derived correlation map should be similar to the ground-truth expectation shown in Supplementary Fig. 2A, right panel (see Methods; Neural data analysis); this happened to be true in the general population (similarity = 0.73, p < 1e-6; Supplementary Fig. 2A). We also found similar results in encode (similarity_encode-suppressive_ = 0.73, p < 1e-6; Supplementary Fig. 2B, left upper panel) and maintenance (similarity_maintenance-suppressive_ = 0.44, p = 0.002; Supplementary Fig. 2B, right upper panel) suppressive as well as enhancive (encode: similarity_encode-enhancive_ = 0.69, p < 1e-5; maintenance: similarity_maintenance-enhancive_ = 0.73, p < 1e-6; Supplementary Fig. 2B, lower panels) neurons. Therefore, there seems to be sophisticated interactions between HPC field activities and the activities of individual neurons; rate and timing of HPC neurons’ spikes with respect to oscillations are seemingly distinct WM mechanisms.

## Discussion

A growing body of evidence suggests that, similar to PFC, HPC neurons change their firing rates during WM^8,14-19^. Here, we find that roughly half the HPC neurons support at least one stage of a verbal WM process with modulations in firing. These neurons are spread across the HPC, existing in both hemispheres and poles. We were then curious to see if the human HPC has functional specializations along the longitudinal axis, similar to the rodent brain, where ventral and dorsal subregions are involved in different behaviors^20,21^. A closer evaluation of the rate-modulated neurons revealed that PH is more robustly reliant on this coding regime. Importantly, this difference is more obvious as the task becomes difficult (greater WM loads). These results are very well-aligned with the existing rodent literature, in which while both subregions are involved in cognition, dorsal HPC has more prominent roles for such behaviors^20,21,27^.

We, and others^16,17^, report that most HPC neurons show tuning to local oscillations, a phenomenon that is stronger in AH. Importantly, AH is also the region with greater tuning to frontal cortex slow oscillations. Thus, we sought to find a mechanism that can possibly mediate these signals. In this line, θ oscillations, which are known as a route for long-range inter-areal communications in the brain^26,29-33^, were synchronized between frontal cortex and HPC. Distant slow oscillation functional connectivity, when happens between frontal and sensory cortices, exerts the top-down cognitive control and tunes the activity of sensory neurons to maintain WM content and/or attend specific information from the incoming inputs^3,9-12,33,34^. Similarly, enhanced frontal cortex-HPC θ coupling in the current study can provide a mechanistic insight on the multicomponent hypothesis of WM, based on which frontal cortex uses storage-related areas, such as HPC, for the maintenance of WM content^16,35-37^. In this case, θ oscillations can act as the coordinating means between frontal cortex and HPC.

Next, we tried to address the relationship between the rate and phase of neuronal firing, which could potentially be a matter of debate. Strikingly, we found that locking to slow oscillations is not correlated with neuronal rate changes. In this line, it has been previously reported that the timing of spikes conveys information that does not exist in the rate of activity; in fact, even the phase of non-rate-modulated neurons’ spikes, with respect to slow oscillations, are informative of the WM content^17^, a finding which further supports the current results. Therefore, since phase and rate modulations are not correlated, we believe that, as suggested earlier^17^, the two mechanisms contain independent information.

In conclusion, we suggest that while both AH and PH have roles in WM processing, neurons in the two subregions behave slightly differently. These results will have the following impacts: First, while there is vast literature on the antero-posterior functional dissociations with respect to behavior in rodents, there is little knowledge in the primate brain. Our findings lay the foundations for further investigations in this direction. Second, we provide additional support for the previous evidence on the cognitive control theory of WM, in which PFC uses storage areas in the brain to maintain information for immediate use. We encourage future studies, especially using more causal methods like perturbation techniques in animals, to provide elaborate details on these ideas.

## Methods

### Dataset

We used a human electrophysiological dataset^25^, which was acquired from 9 drug-resistant epileptic patients (5 female) in 37 recording sessions (one session did not contain oscillatory data). The data contained scalp EEG as well as deep brain neurophysiological recordings from HPC, entorhinal cortex, and amygdala^25^. Of note, the recording sites were solely selected based on the medical diagnostic purposes^25^. For the details of participant information, behavioral findings, as well as the data collection and preprocessing pipelines, see the source publication^25^.

### Behavioral task

In each recording session, the subject performed 50 trials in a modified Sternberg verbal WM task, with the following sequence: 1 sec of fixation, 2 sec of encoding, 3 sec of maintenance, and 2 sec of retrieval (Fig. 1A). During the encoding, 4, 6, or 8 English letters appeared on the screen, during trials of load 1, 2, or 3, respectively, which the subject should have held in mind for the 3-sec period of maintenance; in the retrieval section, the subject should have responded whether the newly presented letter, i.e., probe letter, existed in the encoded items (Fig. 1A).

### Neural data analysis

All neural data analyses were performed in Matlab 2019b or Python v3.8.18. Also, analyses were performed on correct trials, unless noted otherwise. To identify encoder/maintainer neurons, the distribution of time-averaged firing rates of each neuron in the encoding/maintenance section of the trials (n = number of correct trials in the session) were statistically compared to the fixation section using two-sided Wilcoxon signed-rank test in Matlab (with *ranksum* function). We computed the discriminability index (d’; see the Statistical analyses section for mathematical notation), by comparing encode or maintenance firing rate to fixation period, separately for each neuron in each condition. Positive/negative d’ shows more/less activity during the task relative to fixation. Also, higher absolute values of d’ represent greater magnitude of activity changes, regardless of the d’ sign.

SPL was computed using custom written Matlab codes. For locking to the HPC LFP, we first applied wavelet transform between 1-30 HZ (0.5 Hz step) on LFP data. Next, we extracted the instantaneous phases of LFP signals by computing the angles of the analytical signal from the wavelet transform. At the time of each spike, we defined a vector with the simultaneous signal phase and an amplitude of one. All the spike-phases of a neuron in a session were pooled together which were then subject to a vector averaging method, that is unbiassed to sample size, namely pairwise-phase consistency (PPC)^38^, to compute the magnitude of the raw SPL value, through the following:

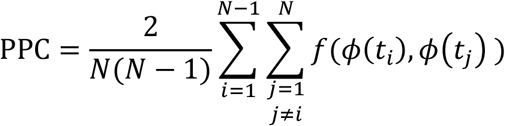

where *N* is total number of spikes and *ϕ*(*t*_*i*_) (or *ϕ*(*t*_*j*_)) shows the signal phase at *i*-th (or *j*-th) spike time and *f* is the function that computes the dot product between two unit-vectors *ω* and *θ* as:

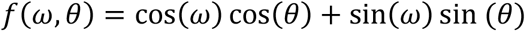

Next, spike times were shuffled across time and trials for 200 times to form the shuffled distribution of SPL values, based on which we computed the shuffle-corrected SPL as the following:

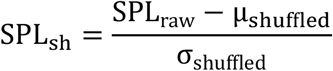

in which μ_shuffled_ and σ_shuffled_ are the mean and standard deviation of the shuffled distribution of SPL values. Also, to find out whether or not the spikes have a tendency towards any specific direction in the polar space, in contrast to being uniformly distribute, we computed the Rayleigh test; a Rayleigh test p-value > 0.05 suggests a uniform distribution of spike-phases in the polar space, which means the SPL values are negligible. At every frequency, the maximal SPL value across all recording sites with significant Rayleigh tests entered subsequent analyses. The preferred frequency of each neuron was defined as the frequency with maximal SPL value in general (i.e., 1-30 Hz, F_overall_) and in θ or αβ bands (F_θ_ and F_αβ_, respectively).

The same process was performed when the signals were EEG, instead of HPC LFP. Of note, to compute the phase locking of HPC neurons to EEG, hemispheric lateralization was respected, i.e., the SPL for neurons in the left (or right) HPC was computed with respect to the F3 and O1 (or F4 and O2) electrodes. Also, since there was only one channel (i.e., recording site) for each location (compared to several sites available in the HPC recordings), it was not required to compute the maximal SPL value at each frequency.

PPL, as a functional connectivity marker, was computed between frontal EEG and HPC LFP. To that aim, we filtered both signals using wavelet transform between 1-30 HZ. After extracting instantaneous phase values of both signals, raw PPL values were computed as the circular average of phase difference between two signals in a trial-wise manner, as the following:

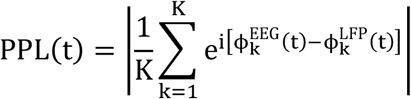

in which 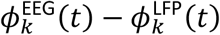 shows phase difference between EEG and LFP signal in trial *K* at time *t*. Z-transformed PPL was calculated the same by z-scoring instantaneous PPL values to the fixation period.

RSA was used to investigate the relation between rate- and phase-coding. For that, we first created correlation maps by computing spearman correlation (using scipy *spearmanr* function) between every pair of metrics; absolute d’ values were used for suppressive units. The similarity of these maps to the ground-truth expectation (Supplementary Fig. 6A, right panel) was then computed using Kendall’s tau correlation (with scipy *kendalltua* function).

## Statistical analyses

Statistical analyses were performed in Python v3.8.18, using SciPy v1.10.1 library, and MATLAB 2019b. Circular data statistics were computed with the MATLAB CircStat toolbox^39^. Details of the statistical tests are described in the appropriate context throughout the manuscript. All permutations were repeated 10001 times. All tests were two-tailed and p-values less than 0.05 were considered as significant. d’ for two data distributions was computed as the following:

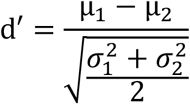

where µ_*i*_ and σ_*i*_ are the mean and the SD of the *distribution*_*i*_ .

## Author Contributions

ZB conceptualized the study. MM and ASM analyzed the data, performed visualizations, and drafted the manuscript. All authors reviewed the manuscript. ZB supervised the study.

## Acknowledgements

The authors appreciate the efforts made to generate this high-quality dataset.

## Data Availability

This study used a publicly available dataset of human neurophysiology^25^. The data can be found at https://doi.gin.g-node.org/10.12751/g-node.d76994/.

## Code Availability

Matlab scripts and functions as well as Python notebooks will be made publicly available at https://github.com/mooziri/Paper_HumanWorkingMemory upon publication of this study.

## Supplementary materials

**Supplementary Figure 1.**
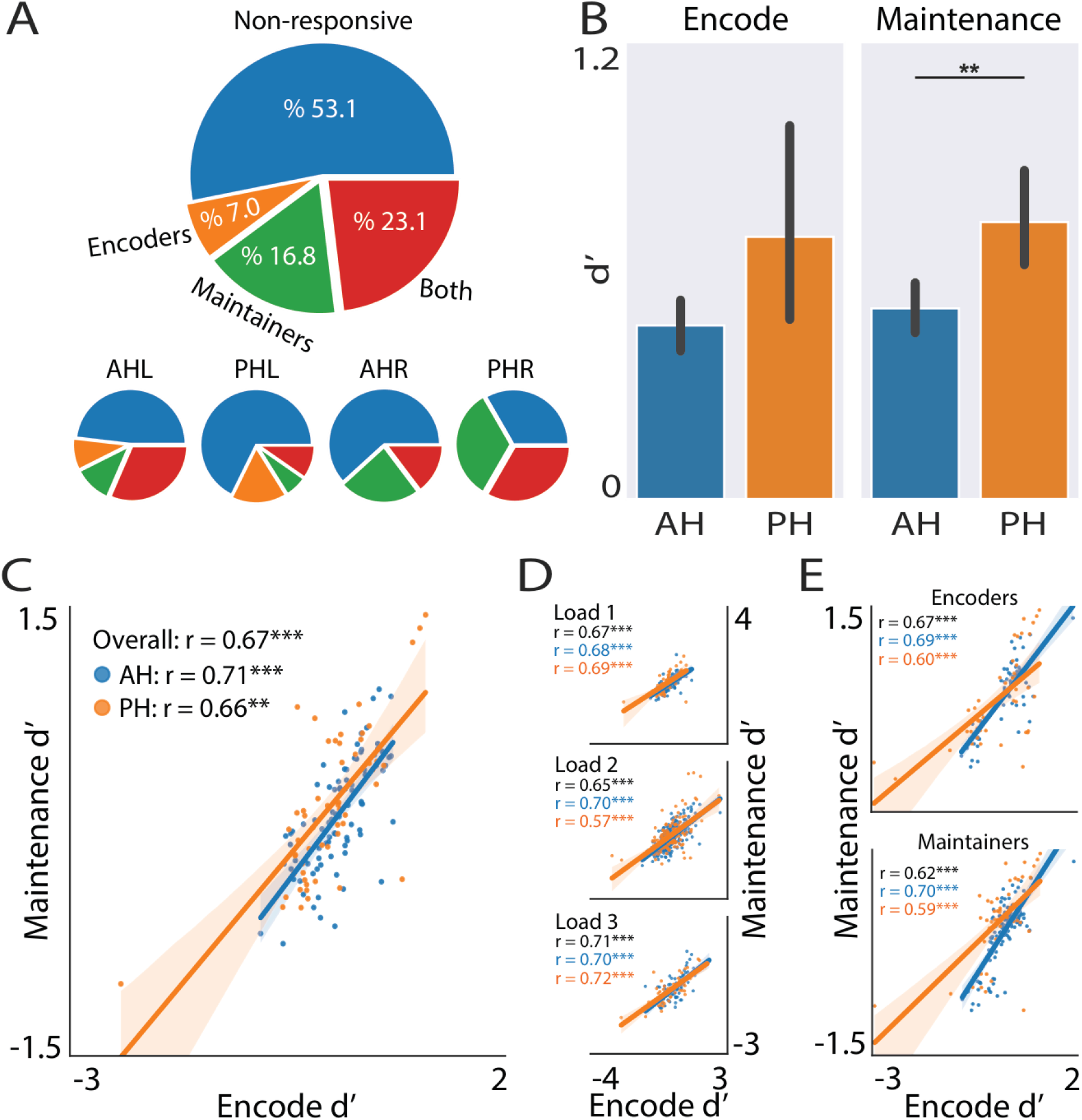
Reproduction of rate-modulation results in the sub-dataset. (A, upper) Pie chart describing the general functional structure of the HPC neuronal population. Assignment of each neuron to the corresponding category was decided based on the results of the two-sided Wilcoxon signed-rank test comparing encoding and maintenance with the fixation firing rates. (A, lower) Same as A, upper, but broken down for each sub-region and hemisphere. (B) Statistical comparison of encode (left panel) and maintenance (right panel) d’ between rate modulated neurons in the AH (n_encode_ = 27, n_maintenance_ = 36; blue columns) and PH (n_encode_ = 16, n_maintenance_ = 21; orange columns). Bars and error bars represent mean and %95 CI, respectively. Mann-Whitney test. (B)Correlation between encode d’ and maintenance d’. Each point is a neuron. Solid lines and shaded areas denote the fitted linear regression models and the %95 CI of the regression. Spearman correlation. (D) Same as C, but for each load. (E) Same as C, but for encoder (upper panel) and maintainer (lower panel) neurons. ** p < 0.01, *** p < 0.001. AH(L), (left) anterior hippocampus; AH(R), (right) anterior hippocampus; PH(L), (left) posterior hippocampus; PH(R), (right) posterior hippocampus.

**Supplementary Figure 2.**
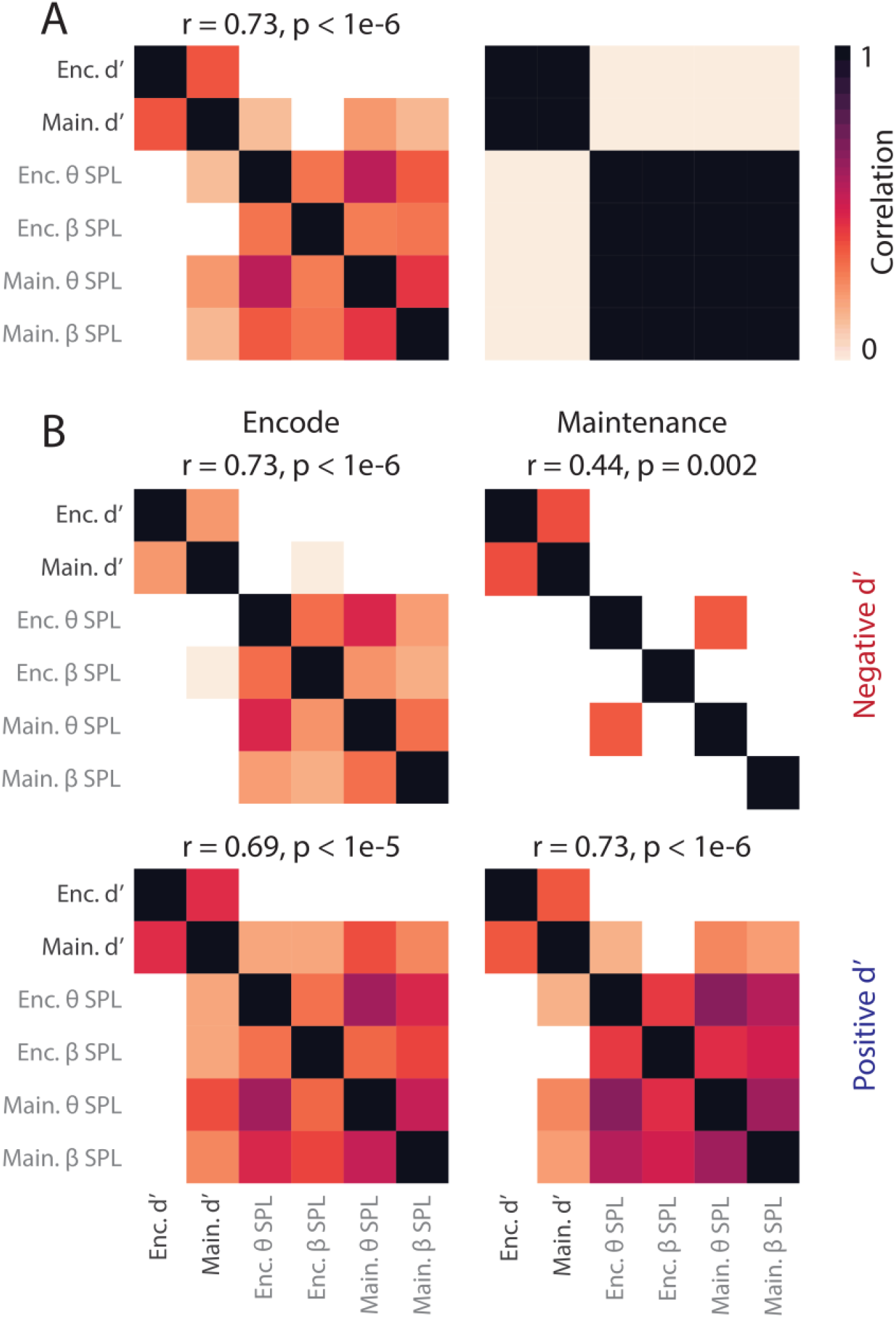
Phase-rate interactions during WM in the HPC approached by RSA. (A, left) Correlation map, created by pairwise Spearman correlation between metrics, in the general HPC population. (A, right) Ground-truth expectation of correlation map, if the two coding regimes are distinct. (B) Correlation maps for encode-suppressive (left upper panel), maintenance-suppressive (right upper panel), encode-enhancive (left lower panel), and maintenance-enhancive (right lower panel) neurons. Similarity values are denoted above each correlation map and were computed by Kendall’s tau correlation. HPC, hippocampus; RSA, representational similarity analysis; SPL, spike-phase locking; WM, working memory.

